# The sex-specific effects of RAGE signaling and Type 2 Diabetes on mouse cortical bone mechanics, structure, and material properties

**DOI:** 10.1101/2025.01.01.631018

**Authors:** Timothy Hung, Kaitlyn S Broz, Remy E Walk, Simon Y Tang

## Abstract

Individuals with type 2 diabetes (T2D) are prone to fracture at numerous skeletal sites despite presenting with a higher bone mineral density (BMD). The accumulation of Advanced Glycation End-products (AGEs) in the bone tissues of patients with T2D could be contributing to this paradox, of increased skeletal fragility with higher BMD. AGEs can impair bone cell homeostasis via the receptor for AGEs (RAGE). To investigate the effects of diabetes, AGE accumulation, and RAGE signaling on mouse cortical bone, we utilized male and female leptin receptor-deficient (db/db) mice from three age groups ranging from 3-14 months of age, which were crossed with animals carrying constitutively RAGE-deficient alleles (RAGE^−/−^). The morphological, mechanical and material outcomes were measured using microCT, 3-pt bending, and an AGE assay. We observed significant impairments dependent on age and sex to the bone matrix and whole-bone mechanical behavior due to diabetes with some impairments alleviated with the ablation of RAGE. In older female diabetic mice, the removal of RAGE signaling prevented the deficits in bone mechanics, morphology and tissue mineral density (TMD). Male diabetic mice without RAGE signaling exhibited improved material properties. The study demonstrated that some bone impairments associated with T2D are prevented with RAGE ablation and may be partially reversible with the inhibition of RAGE signaling.

**Graphical Abstract:** 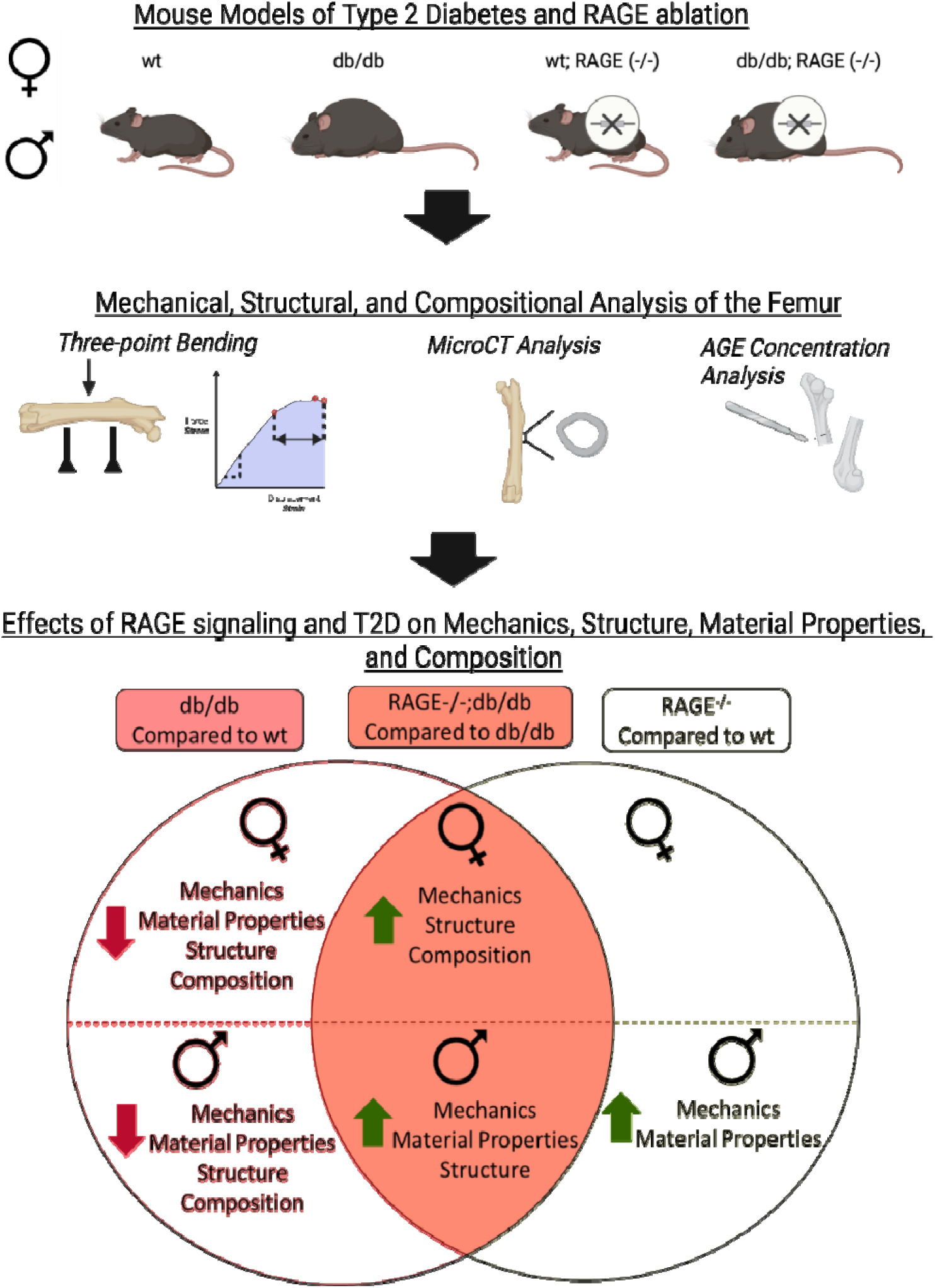

## 3. Introduction

Diabetes is a growing global health concern with its prevalence projected to increase significantly in the foreseeable future.^1^ Among the numerous complications that occur with type 2 diabetes (T2D), the increased incidence of bone fractures is often underappreciated.^2,3^ Patients with T2D suffer more fractures at several skeletal sites despite presenting with a higher bone mineral density (BMD) compared to non-diabetic age-matched individuals.^2,4,5^ This dichotomy makes fractures in patients with T2D difficult to predict and manage. While the clinically utilized fracture risk assessment (FRAX) adjusts for T2D, the adjustment does not appear to adequately represent the fracture incidence in these patients.^6–8^ Identifying the contributors to skeletal fragility in T2D, including matrix level changes, may allow for more effective clinical evaluation of fracture risk in these patients.

The discordance between Dual Energy Xray Absorptiometry (DEXA) and fracture incidence suggests that there are material-level impairments in the bone matrix that contribute to fragility. One possible explanation for this paradox is the accumulation of Advanced Glycation End-Products (AGEs) in the tissues of patients with T2D; which has been implicated in other diabetic complications.^9,10^ AGEs are formed through the non-enzymatic, covalent crosslinking between reducing sugars and amino groups in organ components, such as collagen and other structural proteins in bone.^4^ Chronic hyperglycemia in T2D promotes the formation of AGEs, such as carboxymethyl lysine (CML) and pentosidine. Accumulation of AGEs in the bone matrix negatively impact bone from the molecular level to the overall mechanical performance both *in vitro* and *in vivo*.^11–13^ *In vitro* ribosylation has been used to induce formation of non-enzymatic glycation and AGE-induced crosslinking in bone, but it often results in large, supra-physiological increases in AGE concentrations.^14^ AGE crosslinking in these experiments leads to significant changes in bone mechanics, including post-yield mechanical properties, stiffness, and fracture mechanics.^15–17^ Studies analyzing mechanical properties of human bone tissue from non-diabetic patients and patients with T2D have also shown the deleterious effects of AGEs on bone mechanical behavior.^11^

Utilizing *in vivo* models of T2D may be more physiologically relevant for defining the pathological impact of AGEs on bone. Several mouse models exist that either genocopy or phenocopy genetic and clinical aspects of diabetes^18^, including many models that recapitulate impaired whole-bone fracture resistance, including reduced yield force, stiffness, and ultimate load.^19–21^ Despite T2D having a differential prevalence between males and females^22,23^ and symptoms being exacerbated by aging^24,25^, few studies in preclinical models have included these factors. Therefore, there is a compelling need to include sex and age as variables in investigations of how diabetes affects bone mechanical behavior, material properties, structure, morphology, and tissue composition.

AGEs come as a one-two punch to the musculoskeletal system. AGEs not only directly affect the bone matrix composition and its mechanics, but also matrix modulation by bone cells often in part through activation of the receptor for advanced glycation end-products (RAGE). RAGE signaling stimulates the NF-кB inflammatory pathway and disrupts bone cell homeostasis in a myriad of ways.^26,27^ Inhibiting RAGE in T2D could improve bone cell function and in turn bone mechanics. RAGE knockout mice have enhanced bone cell function, improved bone matrix quality, ^28,29^ and greater whole-bone fracture resistance.^30^ However, these studies have been limited to *in vitro* cell function, male animals, or animals without T2D. Nevertheless, these data indicate that RAGE plays a role in bone cell homeostasis as such we hypothesize that the ablation of RAGE will result in improved mechanics in a sex- and age-dependent fashion in a mouse model of T2D.

## 4. Materials and Methods

### 4.1 Mouse models

All studies were performed with WUSM IACUC approval. One-hundred and sixty one mice were used for this study; mice were distributed across 4 genotypes and 3 age groups (n=6-9 mice/genotype). Mouse genotypes included wild type (wt), RAGE-null (RAGE^−/−^) ^31^, leptin receptor-deficient (db/db)^32^, and double mutant mice (RAGE^−/−^;db/db). All animals were maintained on the C57/BL6J background for least 10 generations prior to entering breeding for this study (Figure 1a).

**Figure 1.**
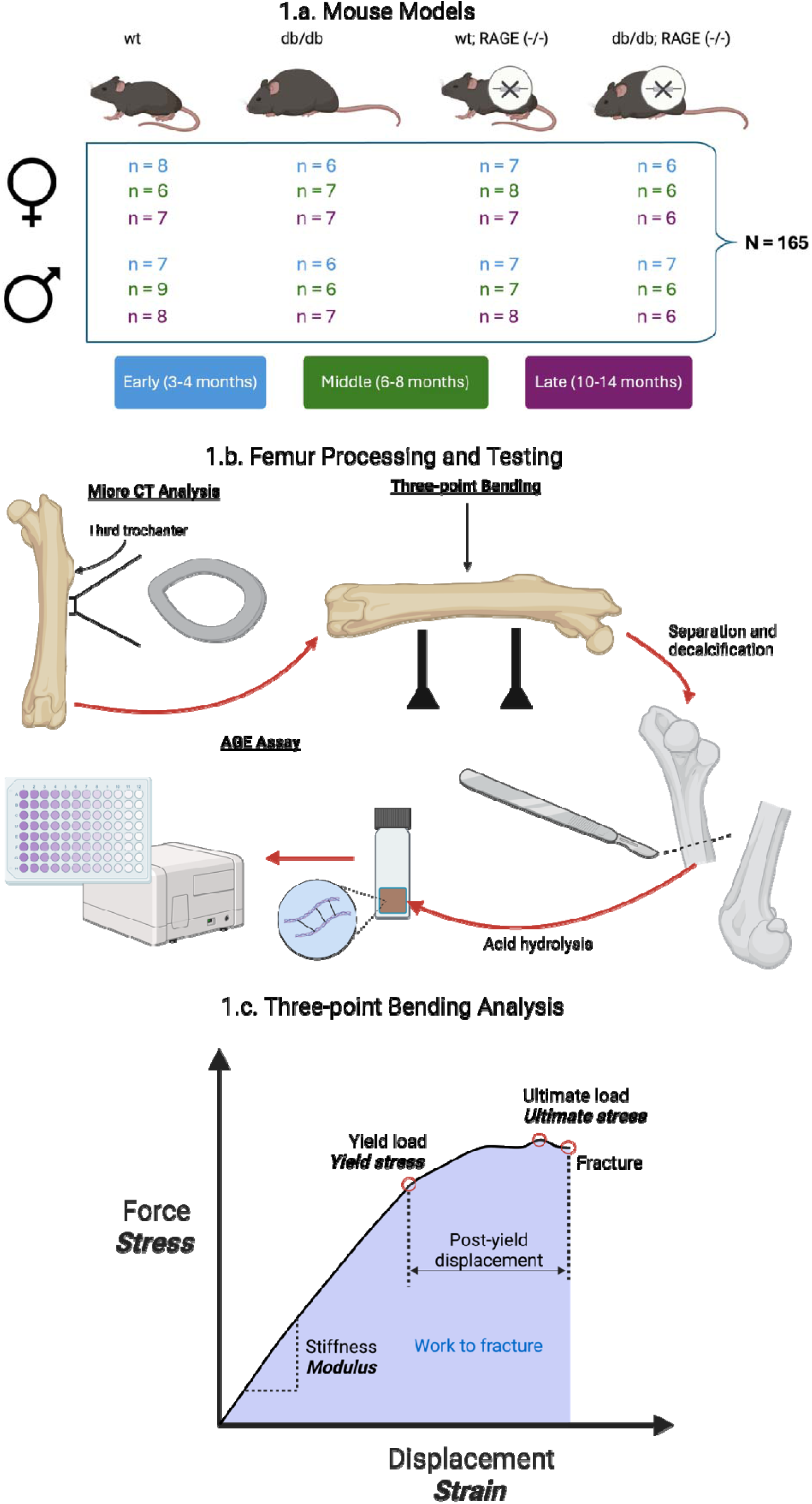
Experimental Design: 1.a Sample sizes for mouse models of T2D and RAGE ablation for Early (3-4 months old), Middle (6-8 months old), and Late (10-14 months old) groups in male and female mice. 1.b Depiction of femur processing and testing. First, microCT was completed and the cortical bone adjacent to the third trochanter was analyzed, followed by three-point bending, decalcification, hydrolysis, and AGE analysis. 1.c. Representative stress-strain curve overlaid on load-displacement curve for three-point bending test. Mechanical and material properties analyzed are indicated on the graph.

Fasting blood glucose was measured using blood drawn from the submandibular vein to confirm diabetic status. The animals were euthanized at one of three time points: 3-4 months (Early), 6-8 months (Middle), and 10-14 months (Late). Following sacrifice, right femurs were dissected, wrapped in PBS-soaked gauze, and stored at −20C. Body mass was recorded at time of euthanasia.

### 4.2 Three-Dimensional Histomorphometry

The femurs were scanned using a Scanco vivaCT40 (Scanco medical AG, VivaCT40, Bassersdorf, Switzerland). MicroCT parameters were a 10.5µm voxel size, 70keV, 170uA, and 300ms integration time. Femurs were thawed for 1 hour in PBS prior to scanning, then potted in agar in batches of 7-11 at a time. For each femur, the first 100 slices distal to the third trochanter were isolated and scanned, and the resulting DICOMs were analyzed using a custom Matlab code.^33^ Cortical thickness (Ct.Th), polar moment of inertia (pMOI), and tissue mineral density (TMD) were recorded.

### 4.3 Mechanical Testing

The femurs were subjected to 3-point bending until fracture using an Instron ElectroPuls E1000 (Instron Electropuls, Norwood, MA, USA). The setup consisted of the anterior surface of the femur facing down, with a span length of 7.5 mm, and a loading rate of 0.1 mm/s. Femurs were thawed for a minimum of 1 hour in PBS prior to mechanical testing. Mechanical properties based on the stress strain curves include stiffness, yield load, ultimate load, post-yield displacement, and work to fracture (Figure 1c). Tissue-level mechanical properties including Young’s modulus, yield stress, and ultimate stress were estimated using morphological parameters from microCT based on beam theory.^34^

### 4.4 AGE assay

After loading the bones to failure in three-point bending, the halves containing the femoral head of each sample were decalcified in Immunocal™ for 3 days and a 2mm section of the diaphyseal region adjacent to the fracture location was isolated for further analysis. Samples were hydrolyzed in 6N HCl at 65°C for 3 hours, pipetted into a 96 well plate in triplicate and then the HCl was allowed to evaporate. Samples were resuspended in PBS for an hour and then imaged for AGE fluorescence (excitation 370nm, emission 440nm) using a BioTek Synergy plate reader. Results were standardized to quinine standards with known concentrations and normalized to the amount of collagen in the bone as measured by hydroxyproline content. Hydroxyproline was quantified using a chloramine-T colorimetric assay standardized to commercially available hydroxyproline standards (BioTek Synergy, absorbance 560nm).^15^

### 4.5 Statistics

Three-way ANOVAs were used to test for the effects of age, diabetes, and RAGE on femoral outcomes were conducted for each sex. After analyzing aging-related trends in the data, separate two-way ANOVAs were run for each time point, again separated by sex. Post-hoc analyses were run as uncorrected Fisher’s LSD (least significant difference) tests between groups when main effects of diabetes, RAGE, or the interaction between diabetes and RAGE were significant. Statistical tests with p-values <0.05 were considered significant for both post-hoc and main effects with adjustments for multiple comparisons. Additionally, a principal component analysis was preformed using the combined data from all groups. All statistical tests were run in GraphPad Prism 10 (GraphPad Software, Boston, Massachusetts USA).

## 5 Results

### 5.1 Loss of RAGE in db/db animals does not protect against the overweight and hyperglycemic phenotype

Both the db/db and RAGE^−/−^;db/db mice had increased body mass compared to their respective controls for both sexes at all time points (S1). Leptin receptor-deficient mice typically become diabetic between 1 and 2 months.^19^ The blood glucose of the db/db and RAGE^−/−^;db/db mice was consistently higher than the wt and RAGE^−/−^ controls (S1). In the Late group, several of the female db/db and RAGE^−/−^;db/db mice had significantly lower blood glucose, likely due to failing health. No exclusions were made based on blood glucose in these data.

### 5.2 Diabetes fosters impairments in bone across multiple length scales

The diabetic phenotype was associated with significant impairments in bone mechanics, morphology, and composition in male and female mice. Cortical thickness is decreased in the db/db mice, with greater losses with age in the female mice. Cortical TMD is also decreased in both sexes in the diabetic mice, with an increasing severity with age in the female mice. AGE concentration is increased in the female db/db mice with no effect of age (S3). Stiffness is reduced in both male and female diabetic mice, with more prominent impairments in female mice at later time points. Maximum load is also decreased in female diabetic animals, but not in male mice. Male mice also exhibited reduced post-yield displacement with diabetes. Overall, there is a sex-dependent effect on bone mechanics in these diabetic mice, with a more severe phenotype in the female mice compared to the male mice. Significant impairments in bone phenotype of female diabetic mice are not detected until 6-8 months; improvements in bone mechanics were actually noted in the db/db group in the early time point (Table 1).

**Table 1.** Female early age group (3-4 months old): Mechanics, Material Properties, Morphology, and Composition. For each group and outcome the mean ± standard deviation is reported, statistics are also denoted in the table; ANOVA effects with values less than 0.1 are included in the table, values less than 0.05 are bolded, significant post-hoc differences between groups are denoted with matching letters (a or b).

| | | Average ± Standard Deviation | | | | ANOVA | | |
| --- | --- | --- | --- | --- | --- | --- | --- | --- |
|  |  | wt | db/db | RAGE<sup>-/-</sup> | RAGE<sup>-/-</sup>;db/db | Diabetes | RAGE | Interaction |
| Mechanical Properties | Stiffness (N/mm) | 73.96 ± 23.44 | 80.58 ± 8.92 | 79.57 ± 17.84 | 87.72 ± 21.31 | | | |
| | Ultimate Load (N) | 13.00 ± 2.15 | 14.26 ± 1.09 | 12.85 ± 1.22 | 14.77 ± 2.72 | **0.0433** | | |
| | Yield Load (N) | 8.64 ± 2.56 | 10.36 ± 1.67 | 9.13 ± 1.42 | 11.53 ± 2.56 | **0.0375** | | |
| | Post Yield Displacement (mm) | 0.90 ± 0.61 | 0.72 ± 0.56 | 0.51 ± 0.34 | 0.47 ± 0.23 | | 0.0949 | |
| | Work to Fracture (N*mm) | 7.58 ± 3.27 | 7.22 ± 3.87 | 5.04 ± 3.19 | 5.84 ± 3.11 | | | |
| Material Properties | Youngs Modulus | 6490 ± 1692 | 7166 ± 895 | 8371 ± 2560 | 8216 ± 2778 | | | |
| | Ultimate Stress (N/mm<sup>2</sup>) | 150.45 ± 14.43 | 168.07 ± 13.25 | 170.50 ± 32.46 | 182.95 ± 42.95 | **0.054** | 0.0716 | |
| | Yield Stress (N/mm<sup>2</sup>) | 99.50 ± 25.84 | 122.43 ± 22.78 | 120.89 ± 27.82 | 142.17 ± 35.29 | | | |
| Morphology | Ct.Th (mm) | 0.1831 ± 0.02 | 0.1923 ± 0.01 | 0.1867 ± 0.03 | 0.1898 ± 0.01 | | | |
| | pMOI | 0.2728 ± 0.05 | 0.2852 ± 0.03 | 0.2530 ± 0.09 | 0.2717 ± 0.06 | | | |
| Composition | AGE/HP (ug/ml)*10<sup>2</sup> | 0.3254 ± 0.22 | 0.5230 ± 0.09 | 0.3248 ± 0.12 | 0.3261 ± 0.17 | | | |
| | TMD (g/cm<sup>3</sup>) | 1068.7 ± 55.57 | 1024.4 ± 22.91 | 1040.9 ± 25.23 | 1053.2 ± 38.07 | | | |

### 5.3 The changes in bone mechanics due to diabetes differed between male and female animals in the early age group

There were significant alterations due to diabetes in both male and female mice, with some minimal effects with RAGE ablation in the early age group (3-4 months old) (Table 1, 2). Female diabetic mice exhibited a significant increase in ultimate load, yield load, and ultimate stress, with no significant post-hoc tests, and no significant effects of RAGE signaling. In the male mice, there was a significant reduction with diabetes in stiffness, post-yield displacement, and work to fracture. Male RAGE^−/−^ mice exhibited significantly greater Young’s modulus compared to the wt group; however, the RAGE^−/−^;db/db group did not show a statistical improvement over the db/db mice. Overall, mechanical alterations were noted in male and female mice in this age group due to diabetes with no significant RAGE^−/−^;db/db interactions.

**Table 2.**
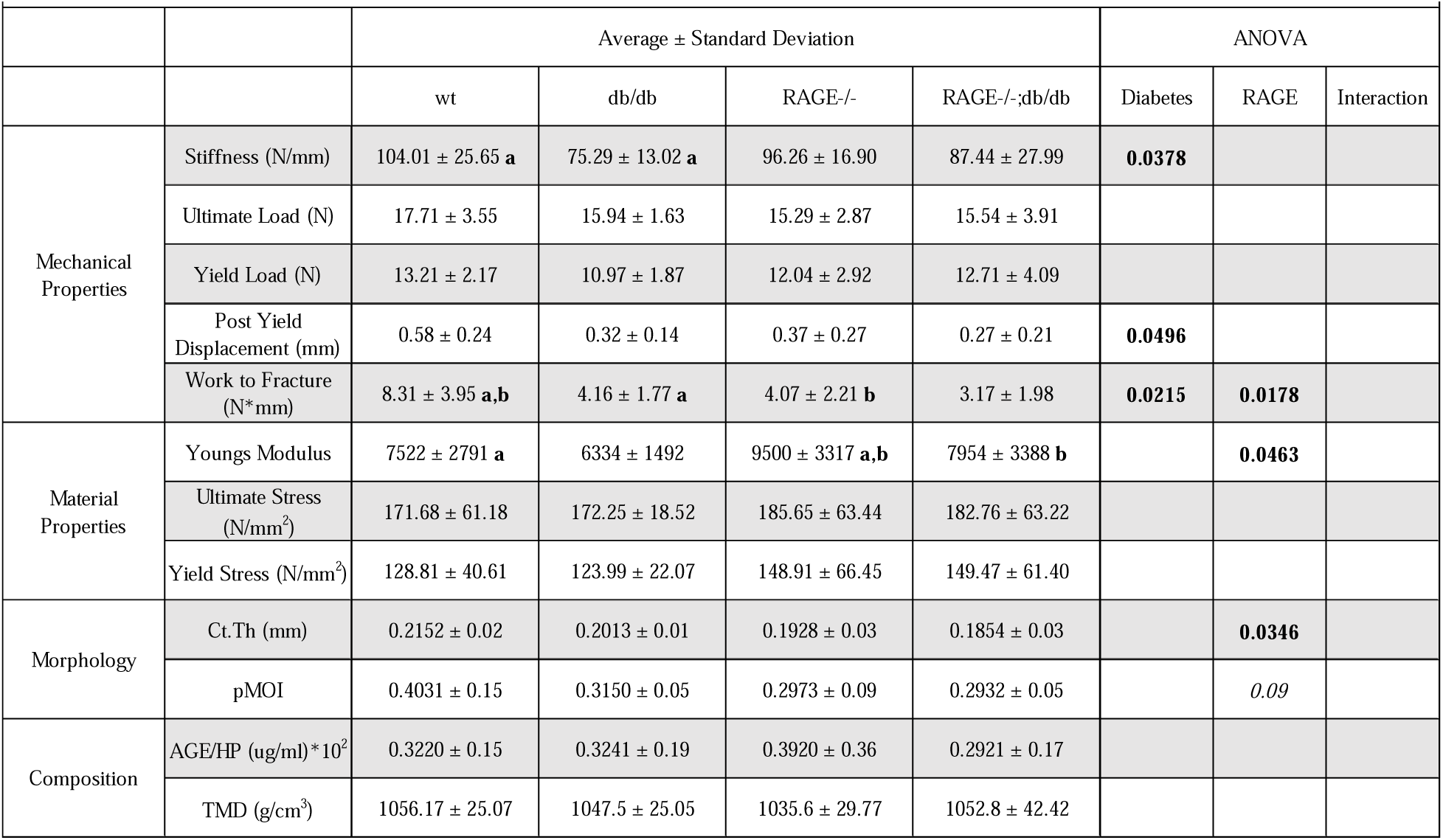
Male early age group (3-4 months old): Mechanics, Material Properties, Morphology, and Composition. For each group and outcome the mean ± standard deviation is reported, statistics are also denoted in the table; ANOVA effects with values less than 0.1 are included in the table, values less than 0.05 are bolded, significant post-hoc differences between groups are denoted with matching letters (a or b).

### 5.4 RAGE ablation mitigated diabetic bone changes in the middle age group for both sexes

Multiple outcomes in the middle age group (6-8 months old) showed significant effects of diabetes, RAGE and the interaction in both sexes (Table 3, 4). The female db/db mice had reduced stiffness, ultimate load, yield load, yield stress, cortical thickness, and TMD with significant post-hoc comparisons between db/db and wt. Some of these deficits were mitigated with the ablation of RAGE signaling, specifically ultimate load, yield load, cortical thickness, and TMD, which all had significant interaction terms. Post-hoc analyses revealed significant improvement in TMD in the RAGE^−/−^;db/db group compared to RAGE^−/−^ group. The male mice showed similar trends in the data. Significant deficits due to diabetes were noted in ultimate load, work to fracture, cortical thickness, and pMOI. In male mice, the RAGE^−/−^;db/db group had improved mechanical, material and morphological properties compared to db/db mice, with significant post-hoc comparisons in yield load, ultimate load, ultimate stress, yield stress, and cortical thickness.

**Table 3.** Female middle age group (6-8 months old): Mechanics, Material Properties, Morphology, and Composition. For each group and outcome the mean ± standard deviation is reported, statistics are also denoted in the table; ANOVA effects with values less than 0.1 are included in the table, values less than 0.05 are bolded, significant post-hoc differences between groups are denoted with matching letters (a or b).

| Table 3. Female middle age group: Mechanics, Material Properties, Morphology and Composition |  |  |  |  |  |  |  |  |
| --- | --- | --- | --- | --- | --- | --- | --- | --- |
| | | Average ± Standard Deviation | | | | ANOVA | | |
|  |  | wt | db/db | RAGE-/- | RAGE-/-;db/db | Diabetes | RAGE | Interaction |
| Mechanical Properties | Stiffness (N/mm) | 118.31 ± 33.87 **a** | 89.34 ± 10.68 **a** | 108.99 ± 12.57 | 103.95 ± 20.88 | **0.0436** | | |
| | Ultimate Load (N) | 19.52 ± 3.61 **a** | 14.51 ± 2.22 **a** | 16.97 ± 1.27 | 16.86 ± 2.16 | **0.0106** | | **0.0137** |
| | Yield Load (N) | 13.13 ± 4.25 **a** | 9.49 ± 1.47 **a** | 11.01 ± 1.24 | 11.60 ± 2.24 | | | **0.037** |
| | Post Yield Displacement (mm) | 0.43 ± 0.16 | 0.31 ± 0.17 | 0.49 ± 0.16 | 0.51 ± 0.24 | | 0.0872 | |
| | Work to Fracture (N*mm) | 6.30 ± 1.75 **a** | 3.55 ± 2.18 **a,b** | 6.79 ± 2.00 | 6.50 ± 1.84 **b** | | **0.0341** | 0.0589 |
| Material Properties | Youngs Modulus | 8232 ± 2493 **a** | 6230 ± 1244 **a** | 9516 ± 653 **b** | 7178 ± 906 **b** | **<0.0001** | | |
| | Ultimate Stress (N/mm<sup>2</sup>) | 183.08 ± 27.22 | 142.43 ± 14.18 | 190.67 ± 18.62 | 158.98 ± 18.35 | **0.0505** | | |
| | Yield Stress (N/mm<sup>2</sup>) | 123.05 ± 38.48 **a** | 94.40 ± 19.81 **a** | 123.39 ± 12.51 **b** | 111.08 ± 29.08 **b** | **<0.0001** | | |
| Morphology | Ct.Th (mm) | 0.2372 ± 0.02 **a** | 0.1893 ± 0.02 **a** | 0.2197 ± 0.02 | 0.2088 ± 0.01 | **0.0002** | | **0.0117** |
| | pMOI | 0.3694 ± 0.07 | 0.3623 ± 0.07 | 0.2882 ± 0.03 | 0.3620 ± 0.06 | | 0.0728 | 0.0753 |
| Composition | AGE/HP (ug/ml)*10<sup>2</sup> | 0.4239 ± 0.21 | 0.6989 ± 0.45 | 0.4366 ± 0.12 | 0.3877 ± 0.11 | | | |
| | TMD (g/cm<sup>3</sup>) | 1102.9 ± 21.54 **a** | 1062.6 ± 25.62 **a,b** | 1071.0 ± 20.57 **c** | 1075.1 ± 16.34 **b,c** | **<0.0001** | | **0.0229** |

**Table 4.** Male middle age group (6-8 months old): Mechanics, Material Properties, Morphology, and Composition. For each group and outcome the mean ± standard deviation is reported, statistics are also denoted in the table; ANOVA effects with values less than 0.1 are included in the table, values less than 0.05 are bolded, significant post-hoc differences between groups are denoted with matching letters (a or b).

| Table 4. Male middle age group: Mechanics, Material Properties, Morphology and Composition |  |  |  |  |  |  |  |  |
| --- | --- | --- | --- | --- | --- | --- | --- | --- |
| | | Average ± Standard Deviation | | | | ANOVA | | |
|  |  | wt | db/db | RAGE<sup>-/-</sup> | RAGE<sup>-/-</sup>;db/db | Diabetes | RAGE | Interaction |
| Mechanical Properties | Stiffness (N/mm) | 122.98 ± 13.95 | 108.39 ± 14.96 | 113.30 ± 15.14 | 112.93 ± 23.14 | | | |
| | Ultimate Load (N) | 18.81 ± 2.84 **a** | 15.69 ± 1.82 **a,b** | 18.01 ± 1.73 | 20.12 ± 1.91 **b** | | **0.0429** | **0.0051** |
| | Yield Load (N) | 12.70 ± 2.22 | 11.22 ± 1.95 **a** | 13.33 ± 1.78 | 14.16 ± 1.55 **a** | | **0.024** | |
| | Post Yield Displacement (mm) | 0.44 ± 0.04 | 0.37 ± 0.17 | 0.42 ± 0.05 | 0.39 ± 0.09 | | | |
| | Work to Fracture (N*mm) | 7.07 ± 1.28 **a** | 4.73 ± 2.57 **a,b** | 6.33 ± 0.84 | 6.72 ± 1.58 **b** | | | **0.0377** |
| Material Properties | Youngs Modulus | 8149 ± 1906 | 7796 ± 1550 | 7575 ± 2014 | 8205 ± 1019 | | | |
| | Ultimate Stress (N/mm<sup>2</sup>) | 170.78 ± 29.41 | 157.41 ± 25.33 **a** | 169.79 ± 22.58 **b** | 200.24 ± 12.83 **a,b** | | **0.0332** | **0.0264** |
| | Yield Stress (N/mm<sup>2</sup>) | 115.07 ± 20.79 | 113.31 ± 26.04 **a** | 126.42 ± 26.74 | 141.55 ± 19.82 **a** | | **0.0368** | |
| Morphology | Ct.Th (mm) | 0.2126 ± 0.01 **a** | 0.1915 ± 0.01 **a,b** | 0.2061 ± 0.02 | 0.2107 ± 0.02 **b** | | | **0.0411** |
| | pMOI | 0.4089 ± 0.07 **a** | 0.3338 ± 0.05 **a** | 0.3919 ± 0.08 | 0.3527 ± 0.04 | **0.0287** | | |
| Composition | AGE/HP (ug/ml)*10<sup>2</sup> | 0.3063 ± 0.08 | 0.3266 ± 0.13 | 0.3393 ± 0.08 | 0.3128 ± 0.06 | | | |
| | TMD (g/cm<sup>3</sup>) | 1072.49 ± 27.58 | 1058.2 ± 26.53 | 1087.9 ± 16.16 | 1077.1 ± 34.82 | | | |

### 5.5 The effects RAGE ablation on bone differed by sex in the late age group

In the late age group (10-12 months old), the female mice had significantly reduced stiffness, ultimate load, cortical thickness, and TMD in the db/db mice compared to wt (Table 5). The RAGE^−/−^;db/db mouse bones exhibited higher stiffness and ultimate load compared to the db/db animals. The male mice exhibited fewer genotype-dependent alterations in the late compared to the middle age group (Table 6). The db/db mice showed impaired yield stress with a trending improvement in the RAGE^−/−^;db/db group. Polar moment of inertia was also significantly impacted by diabetes and loss of RAGE.

**Table 5.**
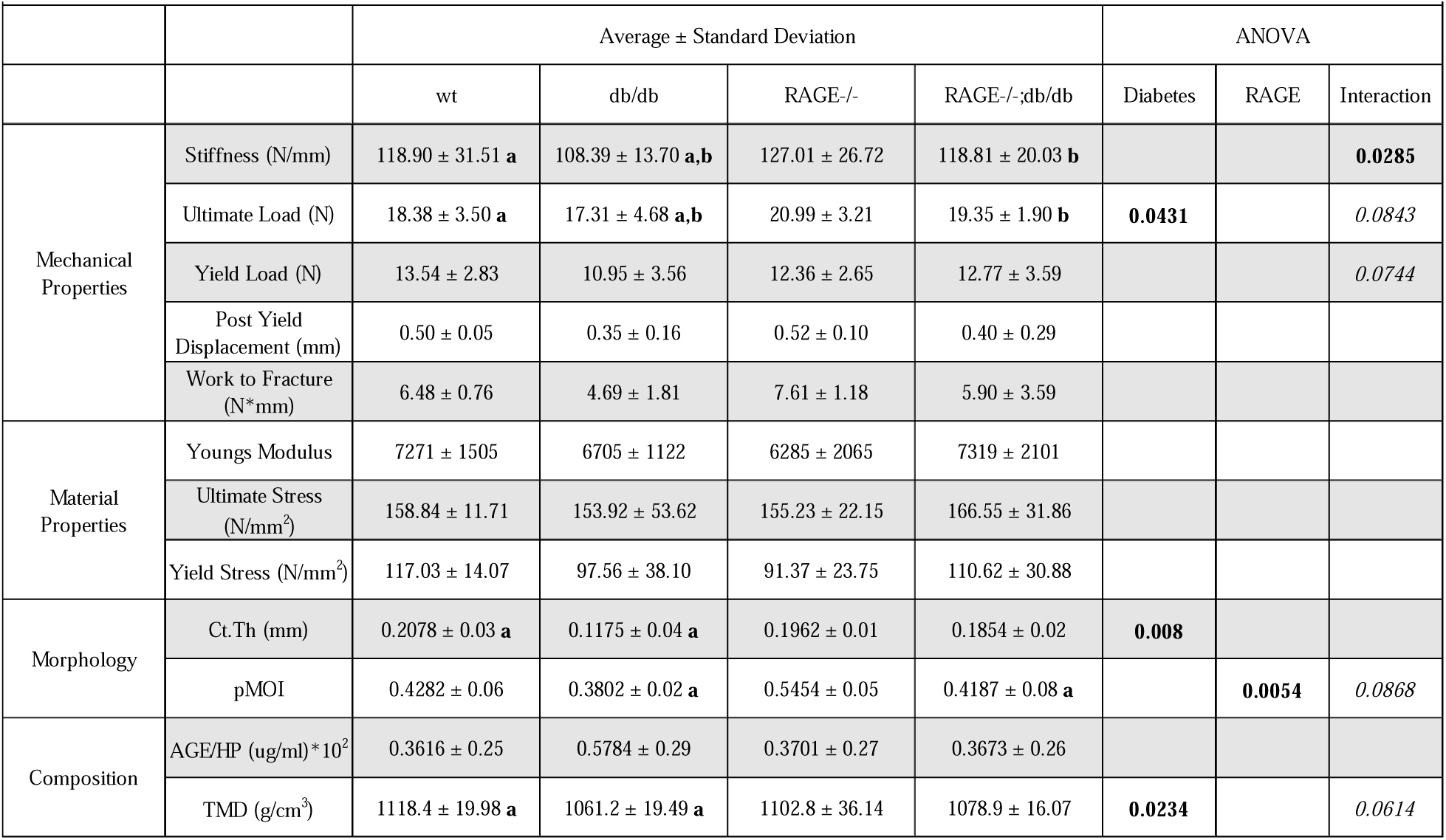
Female late age group (10-12 months old): Mechanics, Material Properties, Morphology, and Composition. For each group and outcome the mean ± standard deviation is reported, statistics are also denoted in the table; ANOVA effects with values less than 0.1 are included in the table, values less than 0.05 are bolded, significant post-hoc differences between groups are denoted with matching letters (a or b).

**Table 6.** Male late age group (10-12 months old): Mechanics, Material Properties, Morphology, and Composition. For each group and outcome the mean ± standard deviation is reported, statistics are also denoted in the table; ANOVA effects with values less than 0.1 are included in the table, values less than 0.05 are bolded, significant post-hoc differences between groups are denoted with matching letters (a or b).

| Table 6. Male late age group: Mechanics, Material Properties, Morphology and Composition |  |  |  |  |  |  |  |  |
| --- | --- | --- | --- | --- | --- | --- | --- | --- |
| | | Average ± Standard Deviation | | | | ANOVA | | |
|  |  | wt | db/db | RAGE-/- | RAGE-/-;db/db | Diabetes | RAGE | Interaction |
| Mechanical Properties | Stiffness (N/mm) | 118.90 ± 20.78 | 108.39 ± 23.52 | 127.01 ± 21.68 | 118.81 ± 15.55 | | | |
| | Ultimate Load (N) | 18.38 ± 3.27 | 17.31 ± 4.70 | 20.99 ± 3.45 | 19.35 ± 4.01 | | | |
| | Yield Load (N) | 13.54 ± 3.06 | 10.95 ± 2.23 | 12.36 ± 3.13 | 12.77 ± 3.04 | | | |
| | Post Yield Displacement (mm) | 0.50 ± 0.27 | 0.35 ± 0.09 | 0.52 ± 0.13 | 0.40 ± 0.13 | | | |
| | Work to Fracture (N*mm) | 6.48 ± 1.63 | 4.69 ± 1.79 | 7.61 ± 2.01 | 5.90 ± 2.72 | **0.0297** | | |
| Material Properties | Youngs Modulus | 7271 ± 1525 | 6705 ± 1115 | 6285 ± 1589 | 7319 ± 1524 | | | |
| | Ultimate Stress (N/mm<sup>2</sup>) | 158.84 ± 25.76 | 153.92 ± 41.01 | 155.23 ± 15.72 | 166.55 ± 46.87 | | | |
| | Yield Stress (N/mm<sup>2</sup>) | 117.03 ± 24.42 **a** | 97.56 ± 19.48 **a** | 91.37 ± 18.87 | 110.62 ± 36.39 | | | **0.0484** |
| Morphology | Ct.Th (mm) | 0.1939 ± 0.02 | 0.1943 ± 0.03 | 0.2045 ± 0.03 | 0.1950 ± 0.01 | | | |
| | pMOI | 0.4282 ± 0.04 **a** | 0.3802 ± 0.05 | 0.5454 ± 0.12 **a,b** | 0.4187 ± 0.05 **b** | **0.0041** | **0.0093** | |
| Composition | AGE/HP (ug/ml)*10<sup>2</sup> | 0.3975 ± 0.20 | 0.4266 ± 0.16 | 0.5448 ± 0.27 | 0.4073 ± 0.14 | | | |
| | TMD (g/cm<sup>3</sup>) | 1102.8626 ± 40.65 | 1071.2869 ± 28.72 | 1118.4513 ± 30.30 | 1081.4520 ± 21.52 | | | |

## 6. Discussion

### 6.1 Severity of diabetic impairments in mechanics increased with age in female mice

The diabetic phenotype significantly impairs bone mechanics, morphology, and composition in male and female mice. Extending the duration of diabetes, increases the severity of impairment in this model, but only in female mice. Material properties of female mice are impaired at the later time points as indicated by reduced Young’s modulus and ultimate stress. Cortical thickness is decreased in the db/db mice in both male and female mice with a greater separation between db/db and wt with aging in female mice only. Cortical TMD is decreased in both sexes due to diabetes, again with an increasing severity with age in the female mice. AGE concentration is increased in the female db/db mice with no effect of age. Overall, there is a sex-dependent effect of type 2 diabetes on bone mechanics in this model, with a more severe phenotype in the female mice compared to male mice. With significant impairments due to diabetes in the female mice not detected until the 6-8 month age group.

The db/db mouse model is a commonly used mouse model of T2D and therefore biomechanics of the mouse bone have been previously characterized^19,35–38^, though most studies tend to focus on one sex and/or one age group. Here we showed that the diabetic phenotype is drastically different in later age groups compared to younger age groups, and the effects vary by sex. At a younger time point, some mechanical deficits have been noted: reduced ultimate load in 11 week old mice^19^ and decreased Young’s modulus in 4 month old mice.^35^ Studies in male mice at older time points, 6-8 months, show similar trends to what we saw in this study, namely decreased ultimate load, yield load, and stiffness^36–38^ or decreased Young’s modulus and increased stiffness and work to fracture^35^, and decreased cortical thickness^38^. Most biomechanical bone studies on db/db mice have been done in male mice only, and to our knowledge there are no comparable femoral biomechanics studies in female db/db mice only.

### 6.2 Diabetes differentially affected the bone mechanical properties between males and females in the between 3-4 month old group

The young male diabetic mice have impaired biomechanics, while younger females have improved biomechanical properties. Interestingly, the diabetic female mice in the early age group have improved mechanical and material properties compared to wt. This phenotype is likely explained by an increase in body mass and the resulting bone adaptations which confer improved biomechanical properties^34^ which occurs prior to the long-term consequences of diabetes that are apparent in the later age groups. The young male mice, on the other hand, had lower stiffness and work to fracture indicating early effects of T2D.

### 6.3 The bone impairments due to diabetes is prevalent in both sexes by 6-8 months of age, and the removal of RAGE protects against multiple bone deficits

In 6-8 month old female mice, diabetes led to impairments in material properties, mechanical properties, morphology, and composition. Ablation of RAGE signaling resulted in improved morphology, mechanics, and TMD. In male mice, impairments due to diabetes were observed in morphology and mechanics, with RAGE^−/−^ animals exhibiting improvements in mechanical and material properties. These data suggest that improvements at the matrix level and morphology in both sexes resulted in improved whole-bone biomechanics. There were no alterations in the RAGE^−/−^ female mice without diabetes, indicating that high-AGE environments are necessary for resultant alterations to the bone matrix mechanics. The male mice, on the other hand, showed improved mechanical and material properties in the animals without diabetes, revealing the sex-specific effects of RAGE signaling. These data align with prior work in male mice.^29,30^ Overall, the bone biomechanical strength and degree of rescue due to RAGE ablation in diabetic animals was similar in the older 10-12 month old mice compared to the 6-8 month old mice. This suggests that extending the duration of diabetes past a certain point, when the effects of diabetes have become saturated, does not lead to further deterioration of femoral mechanics in this mouse model.

### 6.4 Effects of aging and RAGE ablation on bone mechanics vary by sex

In this data set, we were able to identify several aging-related effects on bone mechanics, morphology, composition, and material properties which varied between sexes. Stiffness and ultimate load both increased with age in male and female mice. These data are consistent with what has been reported in mice femoral mechanics^39–41^ and is consistent with the data that shows aging often results in a loss in ductility of the bone matrix. Looking at composition, tissue mineral density increased with age in both male and female mice consistent with what has been shown in C57BL/6 mice previously.^39,42^ Interestingly, AGE concentration was only significantly increased with aging in male mice. AGE concentration has been shown to increase with aging in the bone matrix in both males and female patients with T2D.^43–45^ The accelerated accumulation of AGEs in female db/db animals could be obscuring the changes due to aging alone. Male mice but not female mice experience a significant decrease in material-level mechanics, specifically yield stress, ultimate stress, and Young’s modulus in the 10-12 month age group compared to the 3-4 month group (S1).

Stiffness and ultimate load both increased with age in male and female mice. In female mice, yield load increased with age, while post-yield displacement decreased. Cortical thickness was different in female mice in the 6-8 month age group compared to both the 3-4 and 10-12 month age groups. AGE concentration was significantly increased with age in the male mice. Male mice, but not female mice, exhibit a significant decrease in material level mechanics, specifically yield stress, ultimate stress, and Young’s modulus in the 10-12 month age group compared to the younger groups. AGE concentration in female mice was the only outcome with an overall significant effect of RAGE signaling, where we saw that RAGE^−/−^ animals had reduced AGEs in the bone overall.

The significant statistical interactions between RAGE signaling and age across several outcomes confirms that RAGE ablation exerts its effects in an age-dependent manner. In both male and female mice, loss of RAGE signaling results in a reduced post-yield displacement in only the 3-4 month age group. In the male mice, that trend was also present in the maximum load and cortical thickness, with reduced values in the early age group compared to the older animals.

### 6.5 The decline in femur bone material properties is negatively associated with the accumulation of AGEs

The wide range of AGE concentrations analyzed in this data-set allows for a more thorough analysis to define the associations between AGEs and material, morphological, and mechanical properties. To evaluate the independence of these associations, we performed a principal component analysis (PCA) with the measures pooled from all groups. The PCA revealed that mechanical properties like stiffness and ultimate load group together and are near cortical thickness and tissue mineral density (TMD), suggesting that these structural and material measures are the strongest independent contributors to the mechanical properties in their proximity. Additionally, the material properties, ultimate stress, yield stress, and Young’s modulus are all grouped closely, confirming that these material level modifications and the structural adaptations of bone are similarly affected by diabetes. In the opposite quadrant, independent of the cluster of material properties, the positioning of AGE composition and post-yield displacement indicate that they are characteristics that tend to change in concert (Figure 2).

**Figure 2.**
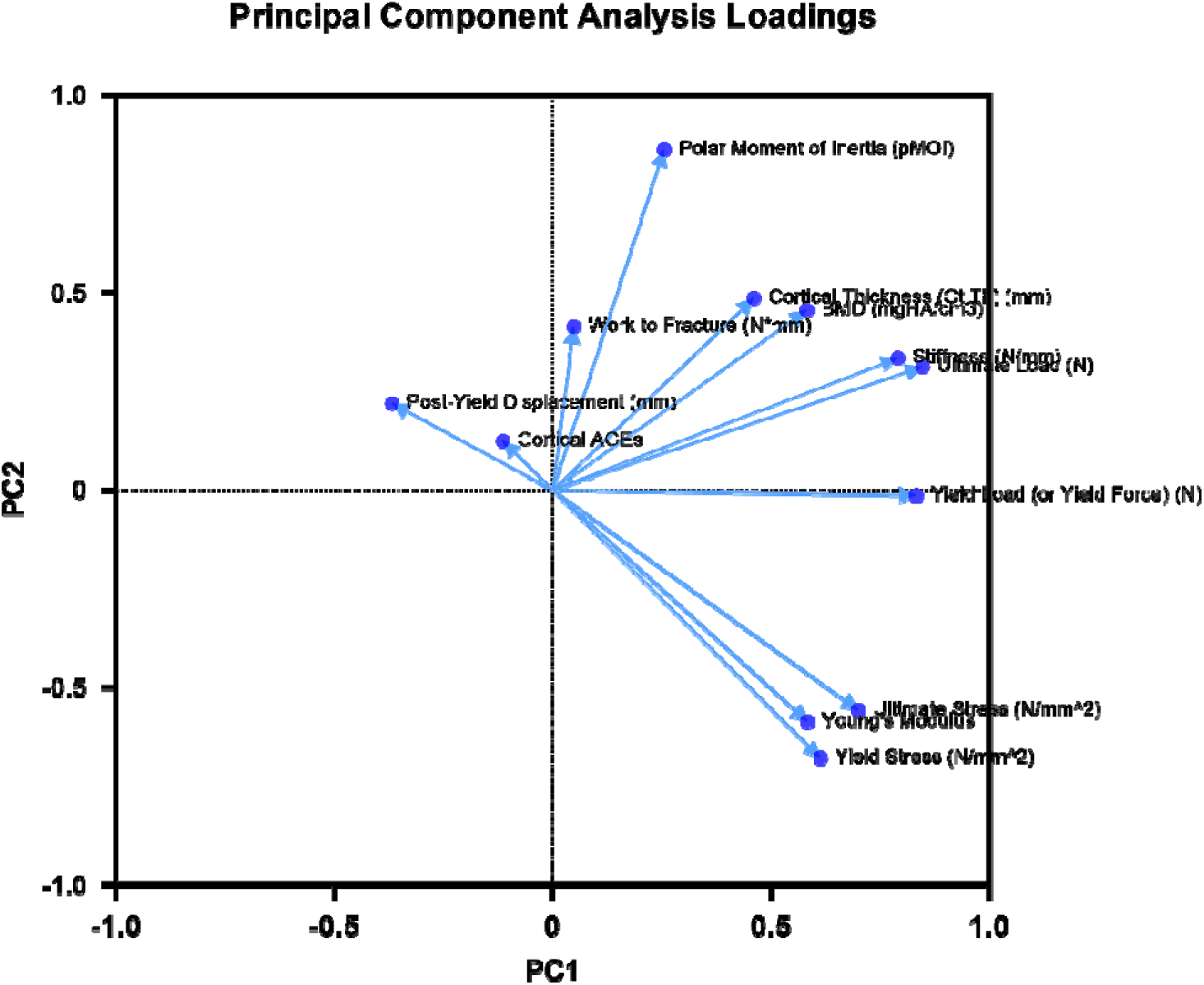
Principal component analysis of multiple measurements across all groups. The PCA revealed that the independence of the various mechanical parameters, as well as the as structural/material measures that are mostly strongly associated with the respective mechanical functions. Further, they are likely governed by similar biological mechanisms, e.g. remodeling. Increased AGEs, due to diabetes, localizes with post-yield displacement. Additionally, the material properties, ultimate stress, yield stress, and young’s modulus are all grouped closely and confirm that these material- and structural-level modifications and the subsequent mechanical behavior of bone are similarly affected by diabetes.

### 6.6 Limitations

There are a few limitations that must be acknowledged in this study. The first is the db/db mouse model of diabetes, which phenocopies the clinical manifestations of obesity, hyperglycemia, and elevated HbA1c through hyperphagic behavior driven by a loss of function mutation in the leptin receptor. Leptin signaling has been shown to orchestrate immune responses^46^, especially during obesity.^47^ Therefore, due to the loss of leptin signaling, these mice likely do not full recapitulate the complex metabolic derangement that occur with prolonged and advanced diabetes. However, since nonenzymatic glycation is a passive biophysical process, the chronic hyperglycemic environment likely continued to enable the formation of AGEs, and likely captures at least some of the biophysical and physiological process that contributes to bone fragility observed in diabetic patients. Another limitation is our use of fluorescent AGEs, which captures a broad population of AGEs, despite numerous and diverse molecular species of AGEs^14^ that may have disparate, individual contributions to bone fragility. More nuance will be needed in the future to distinguish the contributions of individual AGEs. Finally, the systemic ablation of RAGE raises questions about which cell types drive regulation of bone quality in these animals and in patients, although our findings here support future, more in-depth investigations for utilizing RAGE modulation for diabetic bone fragility.

## 7. Conclusions

Our findings suggest that RAGE ablation offers protection against diabetic bone impairments. The presence and extent of these effects vary by age and sex. Older mice with diabetes exhibit more improvement in bone mechanics with RAGE ablation than younger mice, and diabetic female mice without a functioning RAGE receptor generally experience more protection than diabetic male mice. Effects of RAGE ablation on bone mechanics are more pronounced in environments with elevated AGE levels, such as in T2D. In both male and female mice, improvements due to the ablation of RAGE signaling can be attributed to improved morphology and material level properties. Additionally, differences observed in this study due to age and sex in this diabetic phenotype will help inform future studies. This study highlights the contributions of RAGE signaling in changes to the bone matrix in high AGE environments like T2D indicating therapeutic potential of RAGE inhibition for skeletal fragility in patients with T2D.

## Acknowledgements

We would like to acknowledge the Washington University in St Louis Musculoskeletal Research Center for their help and expertise, specifically Michael Brodt and Matthew Silva, PhD. Additionally, we would like to thank Rachana Vaidya, PhD for her insights. This work was conducted with funding support from National Institute of Health: R01AR074441, R01AR077678, R21AR081517, T32DK108742, S10OD028573, and P30AR074992. This work was supported by a Washington University Biology Summer Undergraduate Research Fellowship. The authors are grateful for the generous gift of the RAGE-null mice from Professor Ann Marie Schmidt of New York University

## Author Contributions

TH-Formal analysis, Investigation, Visualization, Writing; KSB-Conceptualization, Formal analysis, Investigation, Methodology, Validation, Visualization, Writing-original draft, Project administration; REW-Investigation, Review, Editing; SYT-Resources, Supervision, Funding, Writing, Review, Editing.

## 8. Supplemental Materials

### S1. Three-way ANOVA p values

| Female Body Mass |  | Male Body Mass |  |
| --- | --- | --- | --- |
| Source of Variation | P value | Source of Variation | P value |
| Age | 0.0049 | Age | <0.0001 |
| RAGE | 0.1691 | RAGE | 0.4215 |
| Diabetes | <0.0001 | Diabetes | <0.0001 |
| Age x RAGE | 0.0133 | Age x RAGE | 0.2789 |
| Age x Diabetes | 0.0653 | Age x Diabetes | 0.0318 |
| RAGE x Diabetes | 0.1667 | RAGE x Diabetes | 0.1215 |
| Age x RAGE x Diabetes | 0.3847 | Age x RAGE x Diabetes | 0.02 |

| Female Blood Glucose |  | Male Blood Glucose |  |
| --- | --- | --- | --- |
| Source of Variation | P value | Source of Variation | P value |
| Age | 0.0244 | Age | 0.0104 |
| RAGE | 0.1464 | RAGE | 0.1553 |
| Diabetes | <0.0001 | Diabetes | <0.0001 |
| Age x RAGE | 0.7639 | Age x RAGE | 0.2543 |
| Age x Diabetes | 0.6208 | Age x Diabetes | 0.0032 |
| RAGE x Diabetes | 0.2836 | RAGE x Diabetes | 0.1092 |
| Age x RAGE x Diabetes | 0.0006 | Age x RAGE x Diabetes | 0.3831 |

| Female Stiffness |  | Male Stiffness |  |
| --- | --- | --- | --- |
| Source of Variation | P value | Source of Variation | P value |
| Age | <0.0001 | Age | <0.0001 |
| RAGE | 0.0568 | RAGE | 0.5025 |
| Diabetes | 0.0759 | Diabetes | 0.0087 |
| Age x RAGE | 0.4012 | Age x RAGE | 0.5404 |
| Age x Diabetes | 0.0847 | Age x Diabetes | 0.5402 |
| RAGE x Diabetes | 0.017 | RAGE x Diabetes | 0.1718 |
| Age x RAGE x Diabetes | 0.2315 | Age x RAGE x Diabetes | 0.7046 |

| Female Maximum Load |  | Male Maximum Load |  |
| --- | --- | --- | --- |
| Source of Variation | P value | Source of Variation | P value |
| Age | <0.0001 | Age | 0.0037 |
| RAGE | 0.2784 | RAGE | 0.1952 |
| Diabetes | 0.0414 | Diabetes | 0.2131 |
| Age x RAGE | 0.3535 | Age x RAGE | 0.0666 |
| Age x Diabetes | 0.0029 | Age x Diabetes | 0.8764 |
| RAGE x Diabetes | 0.0041 | RAGE x Diabetes | 0.114 |
| Age x RAGE x Diabetes | 0.2391 | Age x RAGE x Diabetes | 0.2364 |

| Female Ultimate Stress |  | Male Ultimate Stress |  |
| --- | --- | --- | --- |
| Source of Variation | P value | Source of Variation | P value |
| Age | 0.9661 | Age | 0.0181 |
| RAGE | 0.1135 | RAGE | 0.294 |
| Diabetes | 0.1146 | Diabetes | 0.5091 |
| Age x RAGE | 0.4456 | Age x RAGE | 0.1848 |
| Age x Diabetes | 0.0033 | Age x Diabetes | 0.4998 |
| RAGE x Diabetes | 0.4501 | RAGE x Diabetes | 0.6835 |
| Age x RAGE x Diabetes | 0.6236 | Age x RAGE x Diabetes | 0.0508 |

| Female Young's Modulus |  | Male Young's Modulus |  |
| --- | --- | --- | --- |
| Source of Variation | P value | Source of Variation | P value |
| Age | 0.239 | Age | 0.047 |
| RAGE | 0.1455 | RAGE | 0.2266 |
| Diabetes | 0.0011 | Diabetes | 0.7703 |
| Age x RAGE | 0.5007 | Age x RAGE | 0.2228 |
| Age x Diabetes | 0.0175 | Age x Diabetes | 0.2022 |
| RAGE x Diabetes | 0.9902 | RAGE x Diabetes | 0.8059 |
| Age x RAGE x Diabetes | 0.1554 | Age x RAGE x Diabetes | 0.1405 |

| Female Tissue Mineral Density |  | Male Tissue Mineral Density |  |
| --- | --- | --- | --- |
| Source of Variation | P value | Source of Variation | P value |
| Age | <0.0001 | Age | <0.0001 |
| RAGE | 0.7776 | RAGE | 0.3925 |
| Diabetes | <0.0001 | Diabetes | 0.0999 |
| Age x RAGE | 0.9052 | Age x RAGE | 0.1146 |
| Age x Diabetes | 0.0072 | Age x Diabetes | 0.5479 |
| RAGE x Diabetes | 0.0004 | RAGE x Diabetes | 0.8739 |
| Age x RAGE x Diabetes | 0.9477 | Age x RAGE x Diabetes | 0.5622 |

| Female Ct.Th |  | Male Ct.TH |  |
| --- | --- | --- | --- |
| Source of Variation | P value | Source of Variation | P value |
| Age | <0.0001 | Age | 0.2931 |
| RAGE | 0.9387 | RAGE | 0.596 |
| Diabetes | 0.0006 | Diabetes | 0.0882 |
| Age x RAGE | 0.987 | Age x RAGE | 0.0384 |
| Age x Diabetes | 0.0022 | Age x Diabetes | 0.8525 |
| RAGE x Diabetes | 0.038 | RAGE x Diabetes | 0.411 |
| Age x RAGE x Diabetes | 0.1068 | Age x RAGE x Diabetes | 0.2751 |

| Female AGEs |  | Male AGEs |  |
| --- | --- | --- | --- |
| Source of Variation | P value | Source of Variation | P value |
| Age | 0.2576 | Age | 0.0277 |
| RAGE | 0.0417 | RAGE | 0.4475 |
| Diabetes | 0.0619 | Diabetes | 0.3844 |
| Age x RAGE | 0.9178 | Age x RAGE | 0.8408 |
| Age x Diabetes | 0.9949 | Age x Diabetes | 0.848 |
| RAGE x Diabetes | 0.0314 | RAGE x Diabetes | 0.1975 |
| Age x RAGE x Diabetes | 0.8838 | Age x RAGE x Diabetes | 0.8298 |

| Female Yield Load |  | Male Yield Load |  |
| --- | --- | --- | --- |
| Source of Variation | P value | Source of Variation | P value |
| AGE | 0.0214 | Age | 0.6673 |
| RAGE | 0.2254 | RAGE | 0.1731 |
| Diabetes | 0.4672 | Diabetes | 0.21 |
| AGE x RAGE | 0.647 | Age x RAGE | 0.4869 |
| AGE x Diabetes | 0.0142 | Age x Diabetes | 0.8622 |
| RAGE x Diabetes | 0.0089 | RAGE x Diabetes | 0.0205 |
| AGE x RAGE x Diabetes | 0.3316 | Age x RAGE x Diabetes | 0.9641 |

| Female Yield Stress |  | Male Yield Stress |  |
| --- | --- | --- | --- |
| Source of Variation | P value | Source of Variation | P value |
| Age | 0.38 | Age | 0.0028 |
| RAGE | 0.1776 | RAGE | 0.1245 |
| Diabetes | 0.4565 | Diabetes | 0.8494 |
| Age x RAGE | 0.259 | Age x RAGE | 0.2451 |
| Age x Diabetes | 0.0108 | Age x Diabetes | 0.8908 |
| RAGE x Diabetes | 0.3494 | RAGE x Diabetes | 0.1957 |
| Age x RAGE x Diabetes | 0.7446 | Age x RAGE x Diabetes | 0.6722 |

| Female Post Yield Displacement |  | Male Post Yield Displacement |  |
| --- | --- | --- | --- |
| Source of Variation | P value | Source of Variation | P value |
| Age | 0.0028 | Age | 0.5786 |
| RAGE | 0.4885 | RAGE | 0.472 |
| Diabetes | 0.4194 | Diabetes | 0.0051 |
| Age x RAGE | 0.023 | Age x RAGE | 0.0565 |
| Age x Diabetes | 0.8631 | Age x Diabetes | 0.2369 |
| RAGE x Diabetes | 0.48 | RAGE x Diabetes | 0.5726 |
| Age x RAGE x Diabetes | 0.9198 | Age x RAGE x Diabetes | 0.3208 |

| Female Work to Fracture |  | Male Work to Fracture |  |
| --- | --- | --- | --- |
| Source of Variation | P value | Source of Variation | P value |
| Age | 0.0243 | Age | 0.0471 |
| RAGE | 0.6079 | RAGE | 0.5665 |
| Diabetes | 0.156 | Diabetes | 0.0004 |
| Age x RAGE | 0.0222 | Age x RAGE | 0.003 |
| Age x Diabetes | 0.4236 | Age x Diabetes | 0.412 |
| RAGE x Diabetes | 0.1838 | RAGE x Diabetes | 0.035 |
| Age x RAGE x Diabetes | 0.8403 | Age x RAGE x Diabetes | 0.3366 |

| Female pMOI |  | Male pMOI |  |
| --- | --- | --- | --- |
| Source of Variation | P value | Source of Variation | P value |
| Age | <0.0001 | Age | <0.0001 |
| RAGE | 0.8191 | RAGE | 0.7709 |
| Diabetes | 0.2782 | Diabetes | 0.0004 |
| Age x RAGE | 0.0031 | Age x RAGE | 0.0047 |
| Age x Diabetes | 0.4584 | Age x Diabetes | 0.5921 |
| RAGE x Diabetes | 0.0378 | RAGE x Diabetes | 0.689 |
| Age x RAGE x Diabetes | 0.4213 | Age x RAGE x Diabetes | 0.141 |

### S2. Additional data on the mechanical and material behavior of the mouse bones

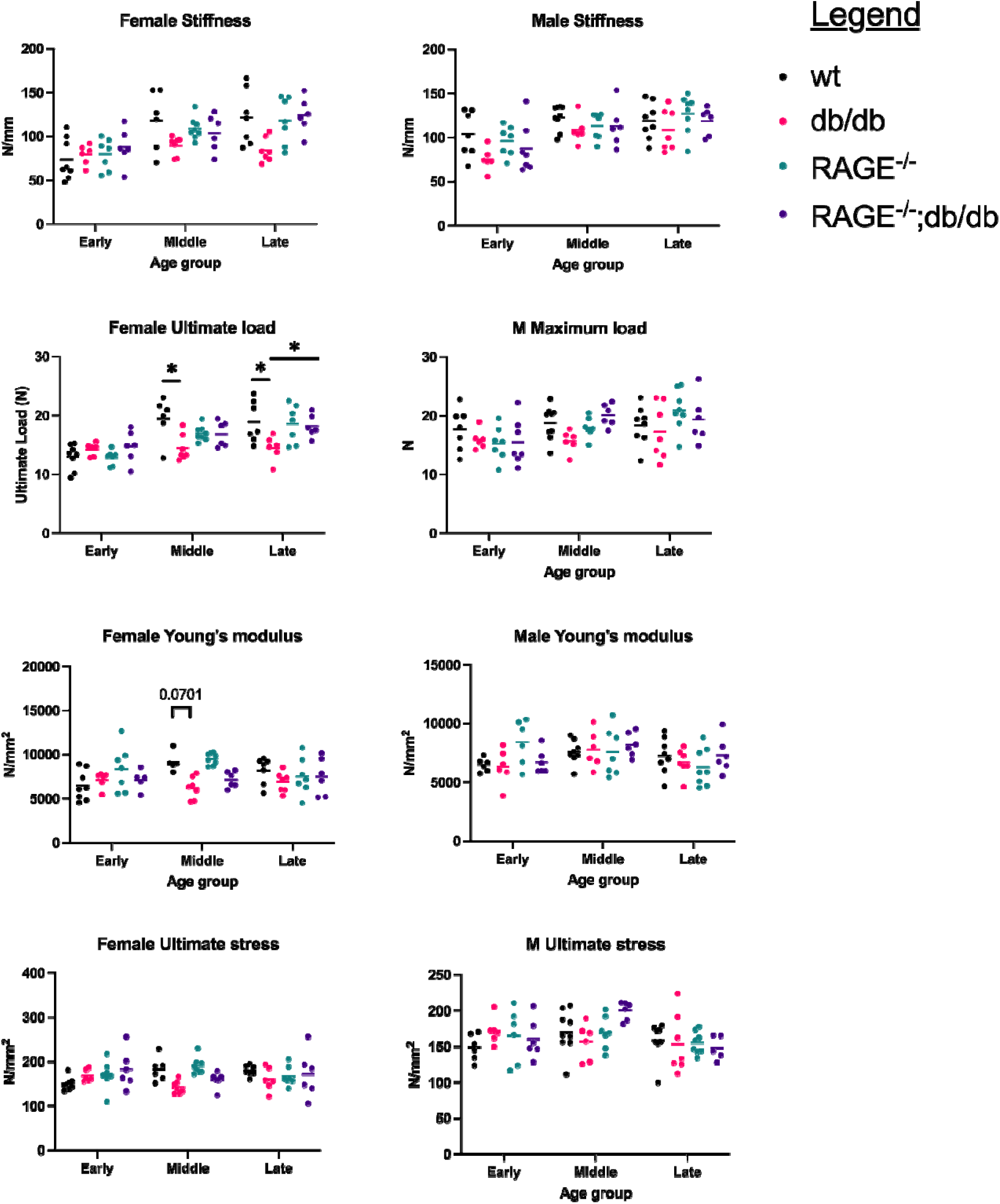

### S3. The age-related changes in bone mechanics vary with sex

**Table 2.1.** The effects of Aging. In female mice there were significant increases in stiffness, ultimate load, yield load, cortical thickness pMOI and TMD due to age, and significantly decreased work to fracture. In male mice stiffness, ultimate load, pMOI, TMD, and AGE concentration increased, and ultimate stress and yield stress significantly decreased due to age.

|  |  | Main Effects of Aging |  |  |  |  |  |
| --- | --- | --- | --- | --- | --- | --- | --- |
|  |  | Female |  |  | Male |  |  |
|  |  | Early | Middle | Late | Early | Middle | Late |
| Mechanical Properties | Stiffness (N/mm) | 80.11<sup>ab</sup> | 105.1<sup>a</sup> | 112.1<sup>b</sup> | 90.75<sup>cd</sup> | 114.4<sup>c</sup> | 118.3<sup>d</sup> |
|  | Ultimate Load (N) | 13.72<sup>ab</sup> | 16.97<sup>a</sup> | 17.55<sup>b</sup> | 16.12<sup>c</sup> | 18.16 | 19.01<sup>c</sup> |
|  | Yield Load (N) | 9.914<sup>a</sup> | 11.31 | 11.91<sup>a</sup> |  |  |  |
|  | Post Yield Displacement (mm)\ | 0.6496<sup>ab</sup> | 0.4359<sup>a</sup> | 0.3587<sup>b</sup> |  |  |  |
|  | Work to Fracture (N*mm) | 6.422<sup>a</sup> | 5.786 | 4.515<sup>a</sup> | 4.925 | 6.212 | 6.168 |
| Material Properties | Youngs Modulus |  |  |  | 6994 | 7794 | 6895 |
|  | Ultimate Stress (N/mm<sup>2</sup>) |  |  |  | 162 | 174.6<sup>c</sup> | 154.1<sup>c</sup> |
|  | Yield Stress (N/mm<sup>2</sup>) | 121.2 | 113 | 111.6 | 137.8<sup>c</sup> | 124.1 | 104.1<sup>c</sup> |
| Morphology | Ct.Th (mm) | 0.188<sup>a</sup> | 0.2138<sup>ab</sup> | 0.1911<sup>b</sup> |  |  |  |
|  | pMOI | 0.2707<sup>ab</sup> | 0.3455<sup>a</sup> | 0.3572<sup>b</sup> | 0.3271<sup>c</sup> | 0.3718<sup>d</sup> | 0.4431<sup>cd</sup> |
| Composition | TMD (g/cm<sup>3</sup>) | 1056<sup>ab</sup> | 1088<sup>a</sup> | 1082<sup>b</sup> | 1051<sup>ab</sup> | 1072<sup>a</sup> | 1088<sup>b</sup> |
|  | AGE (ug/ml)*10<sup>2</sup> | 0.003749 | 0.004868 | 0.004194 | 0.003325 | 0.003213<sup>c</sup> | 0.00444<sup>c</sup> |

## Notes

### Competing Interest Statement

The authors have declared no competing interest.

